# Weight distributions in the fruit-fly and the mouse connectomes

**DOI:** 10.1101/2025.11.10.687553

**Authors:** Michelle T. Cirunay, István Papp, Géza Ódor

## Abstract

By the growing number of available structural connectome data, the distributions of the synaptic weights can be determined which provides a hint at the learning mechanisms at play, both in the global and local level. In this work, we show a numerical analysis of this on the occasion of the latest large and by far, the most well-documented connectomes, the mouse visual cortex and the fruit-fly optical lobe. In literature and in the present work, the synaptic weight distributions for various connectomes follow a power-law (PL) behavior, while the local node strengths can follow heavy-tailed distributions that decay faster. We found that the degree of proofreading on connectomes drastically affects the heavyness of the distribution tails, affecting the interpretation of the structural behavior. In relation to this, there is an ongoing debate on the ubiquitous contradicting observations of lognormal (LN) and PL behavior of weight distributions. Here, we provide an explanation to resolve this by arguing on the basis of generalized central limit theorem. Finally, we show that the global synaptic weight distributions exhibit PL tails with exponents *α* ≥ 3, indicating heavy-tailed, but regular connectivity, while synaptic weights around broadcaster and integrator neurons can be fitted with *α <* 3, i.e have real scale-free fat tails. This suggests a non-random heterogenous organization in which a few dominant synapses facilitate information flow.

**Graphical Abstract:** 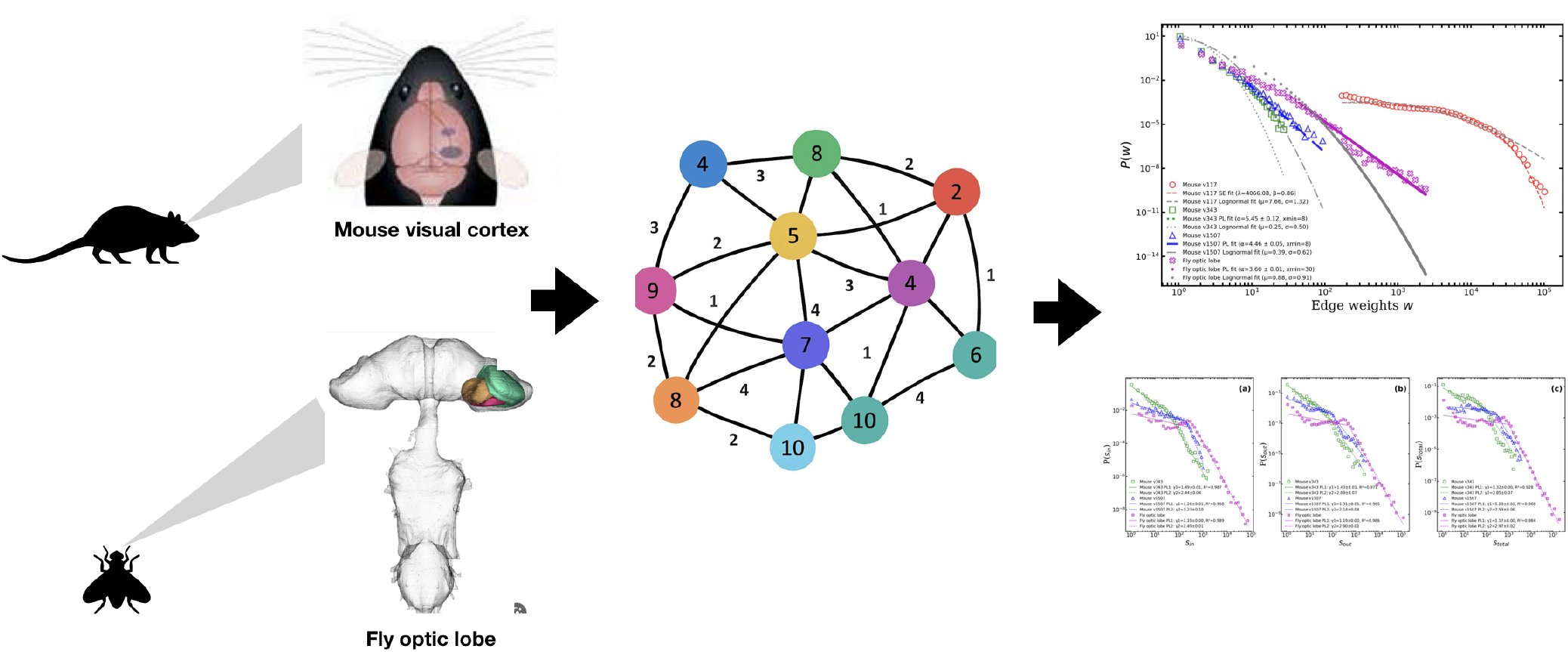

**Highlights:**

- Analyses of the latest large visual connectomes, the mouse visual cortex and male fruit-fly optic lobe, despite being evolutionarily distant showed that their global synaptic weight distributions follow a powerlaw (PL) while their local node strengths follow heavy-tailed behavior that decays faster than a PL providing a hint at the critical learning mechanisms at play
- We briefly illustrate that the degree of proofreading of datasets can affect the heaviness of the weight distribution tails
- The node strength distributions *P* (*s*) for these visual connectomes are found to exhibit dual PL scaling mainly due to the node degree *P* (*k*) heterogeneity with low-*k* nodes contributing to the low-*s* regime, and high-*k* nodes to the heavy tails.
- Edge weight assignments are also found to be non-random and also modulate the behavior of the node strength distributions, although their correlation is below 1%
- We propose an explanation for the deviation of the local behavior from the PL which may resolve the contradicting observation of lognormals and PLs related to critical behavior
- The synaptic weights of links originating and terminating from source and sink nodes are found to also exhibit PL behaviors with exponent less than 3 indicating that they are scale-free

## 1. Introduction

Despite being separated by over 550 million years of evolution, the fly and mouse visual systems have converged on remarkably similar strategies for visual motion processing [1]. The fly’s optic lobe is highly genetically hardwired, featuring a simple, regular columnar structure with invariant connectivity across individuals [2, 3]. In contrast, the mouse visual cortex develops through experience [4, 5] and exhibits hierarchical and modular organization [6, 7], making it a valuable mammalian model for studying visual perception [8], including binocular integration [9]. Both systems process motion using parallel pathways [10], separate ON/OFF channels, and direction-selective neurons namely, T4 and T5 cells in flies [11, 12] and experience-tuned neurons in mouse V1 and higher visual areas [13, 14]. This functional convergence suggests universal computational strategies that transcend species-specific anatomical differences.

Understanding the organizational principles of brain networks remains a central challenge in network neuroscience. The brain criticality hypothesis [15], posits that neural systems operate near a critical point between order and chaos to optimize performance. Evidence from various species supports this idea. For instance, studies in turtles [16] that visual cortex activity becomes more critical, exhibiting scale-invariant dynamics, after exposure to strong sensory input. In mice, recent work [17] shows that during active visual task performance, neural activity in the visual cortex displays signatures of criticality, such as non-Gaussian, scale-invariant patterns identified using renormalization group-inspired methods. These critical states were linked to improved behavioral performance, suggesting that sensory input and learning dynamically tune neural circuits toward criticality.

It is generally known that heavy-tailed statistical distributions are produced by many features of brain dynamics. Prior research has shown log-normal (LN) distributions in the firing rates as well as in the structural connection strengths and suggested generative processes for their phenomena [18, 19, 20]. However, several studies find power law (PL) distributions in brain activity, which are frequently thought to be a result of brain criticality [21, 22, 23]. The physical structure of the connectome has also been reported to exhibit these heavy-tailed statistics recently [24, 25, 26]. These studies focus on optimization logic rather than assuming the backward link of function to structure, incorporating the brain criticality hypothesis. A basic growth model based on mean-field approximations with preferential mechanisms of synaptic re-arrangements, for instance, explains the occurrence of heavy-tailed regimes in the edge weight *P* (*w*) and degree *P* (*k*) distributions of different animal connectomes that have been fitted by PL at the tails [26]. PL tails have been observed in the global weight distributions of the human white matter [27] and fruit fly brain neurons [28]. This PL tail observation for global weights has been reinforced by a more thorough network analysis that includes more complete or partial brain networks [29]. However, it has also offered a distinction with node strength (degree) distributions, which are found to be best fitted by LN functions. Recent experimental and theoretical findings, show that the relation of the structure and function is the strongest in case of criticality [30, 31].

A key mechanism underlying the refinement of these motion-processing circuits is Hebbian learning, which strengthens connections between neurons that are co-activated [32, 33]. In flies, motion detection is largely innate, but early visual experience, particularly within a critical 12–24 hour window after emergence, can influence the development of the optic lobe, notably the growth of photoreceptor terminals (R1–R6) in the lamina. This suggests that even genetically hardwired systems undergo experience-dependent tuning via Hebbian-like mechanisms [34]. In mice, Hebbian plasticity plays a more prominent role in shaping direction selectivity and receptive field organization [35, 36, 37]. However, recent findings also show that co-activity alone does not guarantee synaptic strengthening e.g. neurons with different projection targets may not “wire together” despite firing together [38]. Hebbian plasticity may thus serve as a self-organizing mechanism that pushes both hardwired and plastic systems toward this optimal regime, and in general, can be a plausible mechanism that gives rise to near-critical network organization.

While the aforementioned studies indeed provide evidence of criticality in the brain, we also acknowledge that the presence of heavy-tailed behaviors or similar statistical patterns can arise from a wide class of non-biological or purely topological models. Therefore, rather than simply interpreting scaling behavior as direct evidence of Hebbian learning or critical dynamics, it is necessary to identify more specific structural signatures that are consistent with, and potentially discriminative of, such mechanisms.

In this work, we focus on the joint organization of network topology and edge weights. If activity-dependent processes such as Hebbian learning mechanisms contribute to network formation, we expect (i) a strong coupling between node degree and node strength, reflecting the accumulation of weights on highly connected nodes [39, 40, 41, 42], and (ii) a non-random alignment between edge weights and network structure, such that highly connected regions preferentially support stronger interactions [41, 43, 44]. Importantly, (iii) these features should be sensitive to perturbations that disrupt the relationship between topology and weights [42]. To test these hypotheses, we randomly shuffle a fraction of edge weights while preserving the underlying topology of the networks. This procedure allows us to isolate the role of weight-topology alignment in shaping node-level statistics. If the observed heavy-tailed strength distributions arise purely from topology or random weight assignment, they should remain stable under partial randomization. Conversely, if they depend on structured coupling between weights and connectivity, we expect measurable deviations under perturbation.

By combining distributional analysis with controlled weight randomization and correlation structure measurements, we assess the extent to which empirical network organization is consistent with activity-dependent, self-organizing principles. While our approach does not directly model learning dynamics, it provides a set of falsifiable, structural tests that bridge statistical observations in connectomes with hypotheses about their underlying generative mechanisms.

## 2. Methods and Datasets

### 2.1. Data Sources

In this work, we investigate the properties of the visual systems of the mouse and the fly, which have independently evolved over millions of years but exhibit convergent strategies for motion processing. All previous systematic EM studies of Drosophila visual neurons were based on female brains. Here, we used the connectome of a male fly optic lobe (v1.1), obtained following [45], which can be downloaded via the neuPrint+ client from [46].The fly visual system comprises several anatomically distinct regions: the lamina, medulla, accessory medulla, lobula, and lobula plate. Together, these regions form the optic lobe. However, this particular dataset contains all optic lobe neuropils except for the most peripheral neuropil, the lamina. The entire central nervous system, including the ventral nerve cord, optic lobes, and central brain, is contained in an EM volume that includes this optic lobe, which was the first proofread and examined brain region. This volume contains more than 53,000 unique neurons spanning 732 different types [45].

We also considered the mouse visual cortex downloaded from the MICrONS Explorer static repositories using `minnie65_public`[47]. This dataset includes electron microscopy (EM) image data, segmentations, and their corresponding neuronal meshes for the visual cortex. The dataset covers a volume of 1.4 mm × 0.87 mm × 0.84 mm from a P87 mouse. To illustrate the effect of proofreading and dataset evolution, we consider three versions of the mouse visual cortex: a subset of v117 (the full v117 synapse table is 48GB) with 18,945 nodes and 588,931 edges v343 with 6,261 neurons and 115,379 edges, and (iii) the latest version v1507 with 2,178 neurons and 149,360 edges. Details of these networks are summarized in Table 1. For both v117 and v1507 connectome extraction, we used Python 3.6.8 [48] along with several scientific libraries: NumPy 1.19.5 [49] for numerical computations and array manipulations, SciPy 1.5.4 [50] for signal processing and statistical analysis, and pandas 1.1.5 for structured data handling and analysis.

**Table 1:** Connectome data description.

| Dataset | No. Nodes | No. Edges | Nodes | Edges | Edge Weights |
| --- | --- | --- | --- | --- | --- |
| Male fly optic lobe | 55,571 | 6,533,913 | neurons | synapses | no. synaptic links |
| Mouse visual cortex |  |  |  |  |  |
| v117 | 18,945 | 588,931 | neurons | synapses | no. synaptic links |
| v343 | 6,261 | 115,379 | neurons | synapses | no. synaptic links |
| v1507 | 2,178 | 149,360 | neurons | synapses | no. synaptic links |

To obtain v117, DuckDB 0.3.0 [51] was employed for efficient local querying and joining of large tabular datasets. Neuron identifiers were obtained using CaveClient 5.14.0 [52] from the MICrONS database, specifically from the table `neuron_detection_nv0` by selecting entries with the key `pt_root_id` corresponding to neurons. These identifiers were then cross-referenced with the synapse dataset `synapses_pni_2`, retaining only rows where both `pre_seg_id` and `post_seg_id` appeared among the detected neurons. In sum, this dataset includes all automatically detected neurons (`pt_root_id`) from `neuron_detection_nv0` and all synapses between them from `synapses_pni_2`, without filtering for manual proofreading or verification. On the other hand, we also considered the Mouse visual cortex v343 (released in 2022 February [53]) which have been previously used [24, 25]. This dataset was obtained upon request from the authors and the details of data acquisition can be found in the methods sections of the aforementioned works. Finally, we obtained the most recent materialization v1507, a high-confidence subset of the MICrONS connectome by selecting neurons that had undergone manual proofreading of either axons or dendrites. The proofreading status table (`proofreading_status_and_strategy`) was queried from the public datastack (`minnie65_public`) using the CAVEclient API. Neurons with verified axon or dendrite morphology were extracted, and their unique identifiers `pt_root_id` were used to retrieve synapses. Synapses were filtered to include only connections where both the presynaptic and postsynaptic neurons were proofread, and autapses were excluded. The resulting edge list represents verified synaptic connections between proofread neurons and serves as the basis for subsequent network analyses.

To summarize, we considered three levels of neuron proofreading in the MICrONS dataset: a subset of the raw dataset of all automatically detected neurons and synapses for maximal coverage, a semi-proofread dataset including edges where either neuron is manually verified to balance confidence and completeness, and a fully proofread dataset containing only edges between manually verified neurons for high-confidence connectivity analyses.

### 2.2. Network metrics

We consider weighted and directed networks of the mouse visual cortex and the optic lobe of a male fruit fly. In order to quantitatively compare these two visual systems in consideration, we employ complex network analyses. Here, every node corresponds to a neuron and the synapses are represented by edges. The simplest and most basic quantities are the edge weights *w*_*ij*_, which can describe the multiplicity of synapses between neurons. The measure of how well-connected a node is to its neighbor is determined by the quantity, called the node strength *s*_*i*_, which is the sum of the weights of all edges connected to node *i*,

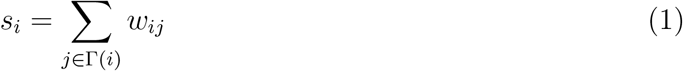

where *w*_*ij*_ is the weight of the edge between nodes *i* and *j*, and Γ(*i*) are the neighbors of node *i*. For directed networks, this splits into in-strength (sum of incoming edge weights) 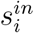 and out-strength (sum of outgoing edge weights) 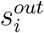.

### 2.3. Null Models

To understand whether the observed dPL behavior arises from the underlying network structure or from edge weight heterogeneity, we construct a set of null models that isolate the roles of network topology and edge weight assignment.

#### 2.3.1. Random Topology Null Model

To assess the role of network topology independent of structural constraints, we construct a random topology null model by generating a random directed graph with *N* and *E* corresponding to the Mouse v1507 for computational ease. Here, all higher-order structures such as the empirical degree sequence, degree correlations, and network motifs are not preserved. As such, this null model serves as a baseline for evaluating whether the observed structural patterns arise solely from network size and density.

#### 2.3.2. Degree-Preserving Null Model

On the other hand, to isolate the effect of the empirical degree structure, we construct a degree-preserving null model, where the exact in- and out-degree sequences of the empirical network Mouse v1507 such that the dPL structure is maintained. Doing so destroys correlations: *i* is assigned with 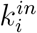 random incoming and 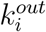 outgoing stubs, and these stubs are randomly matched to form directed edges.

#### 2.3.3. Weight Assignment Schemes

For all null models, the edge weights were assigned independently of the topology, drawn from uniform random (*P* (*w*_*ij*_) ∈ *U* (*w*_*min*_, *w*_*max*_))), or from power-law (PL) distributions 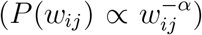. In both cases, the weights were constrained within the same minimum and maximum (*w*_*min*_, *w*_*max*_) as observed in empirical network. For the PL weights, we used the empirical exponent: *α* ≃ 4.46(5) of the synaptic weights, determined by fitting. By using the same range, we ensure that the scale of node strengths is comparable across null models and the empirical network, preventing artificial inflation or suppression of heavy tails. By matching both the exponent and weight ranges allow us to detect differences in *P* (*s*), that can be attributed primarily to topology and weight heterogeneity, instead of the scaling choices.

## 3. Results

Figure 1 shows the global edge weights distributions *P* (*w*_*ij*_) of the datasets being considered. Here, the effect of the degree of proofreading is apparent in the mouse visual cortex datasets. For example, for the Mouse v117 which is a subset of the synapse table first released by the consortium, we observe the edge weights to follow a stretched exponential (SE). In an earlier attempt to mine this dataset, we have started from a very large data frame of more than 300M synapses. Our initial analyses revealed that the weight distribution tails always decay fast, best described by an exponential. Further filtering the synapse table for *purely neurons* led us to this subset of synapses (here we considered as Mouse v117), we observed its tails to have become slightly fatter evolving into a stretched exponential. This observed difference may be attributed to the fact that different analysis pipelines impact the characterization of connectomes [54] i.e. the skewness, right-tailedness are affected. Therefore, the fidelity of connectome mapping methods evolves in time, especially the proofreading, which is currently done mainly manually, thus it proceeds slowly, which can lead to inconsistencies. Here we tried to point out ambiguities coming from data filtering in case of the mouse visual cortex and tried to rely on the most precise proof-read parts in our network analysis.

**Figure 1:**
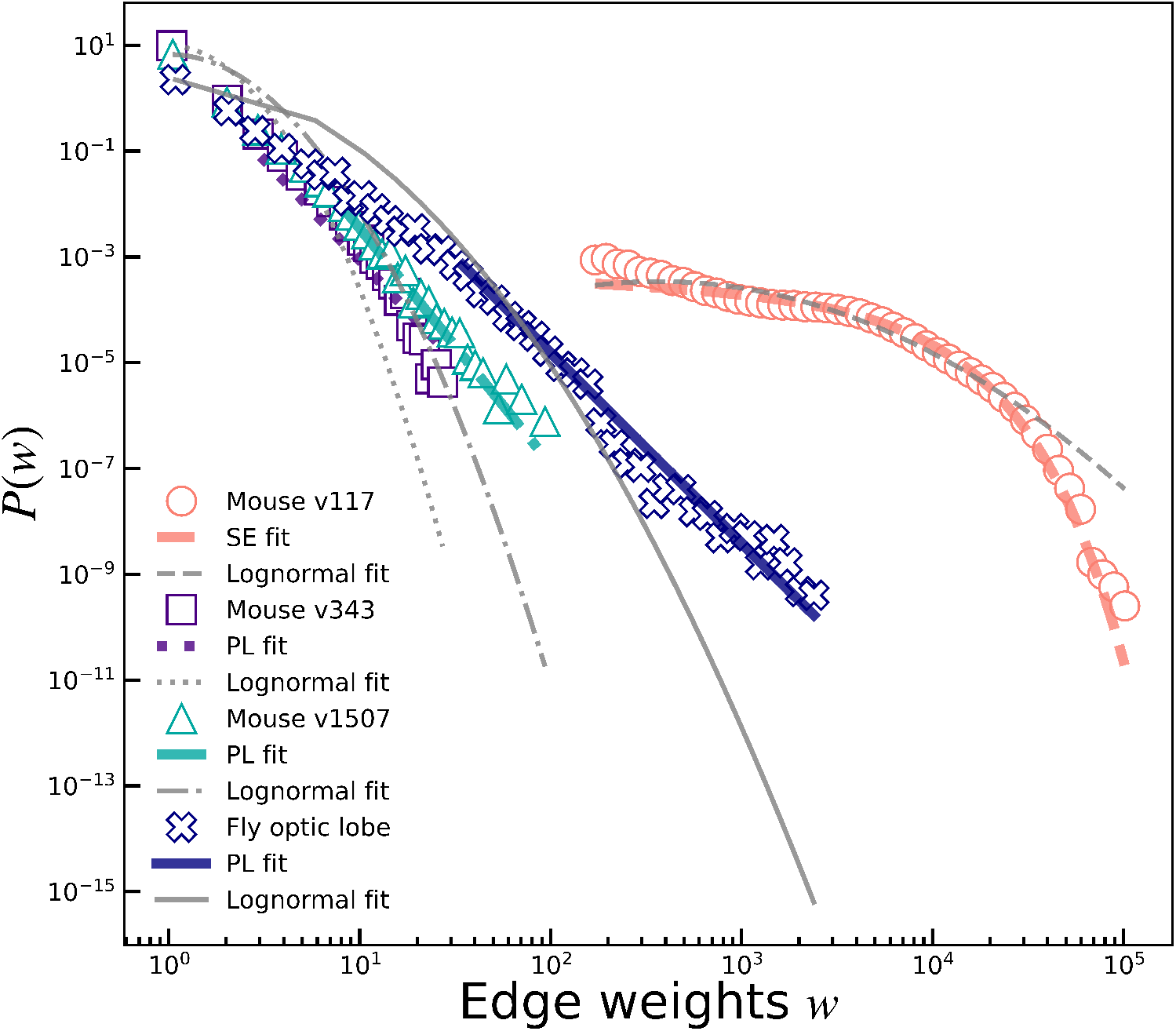
Global edge weight distributions and curve fits. The weight distributions of the *proofread* visual connectomes (Mouse v1507 and Fly optic lobe) still follow a PL behavior, albeit with steeper slopes (*α >* 3). To illustrate the effect of the lack of proofreading, Mouse v117 is included and is shown to behave most likely as a stretched exponential.

Our supposition that the filtering and proofreading lead to fatter tails is supported by the behavior of the tails of Mouse v343 and v1507. While the tails of v343 and v1507 are shorter than three decades to fit a PL, we suppose that subsequent release of proofread mouse cortex data should validate the correctness of this choice of fitting. Here, we proposed some distributions that will most likely fit the behavior of the tails by calculating the Kolmogorov-Smirnov (KS) distance and model selection metrics, the Akaike Information Criterion (AIC) and the Bayesian Information Criterion (BIC) metrics [see Supplementary Material].

In Table 2, we present the KS, AIC, and BIC values for our global edge weight models. For the Mouse v117, the stretched exponential (SE) has a smaller KS (0.3045) than the LN model (0.3615), suggesting that SE captures the bulk and tail behavior better. The same can be observed for the newer mouse cortex versions, v343 and v1507. Meanwhile, for the Fly optic lobe, the PL model has a lower KS (0.3182) than LN (0.3892), suggesting that a PL distribution better describes the network. We use the KS distance for model selection because it directly measures how far the fitted distribution deviates from the data across the full range (see Eqn. C.14), making it sensitive to shape differences. In contrast, AIC and BIC depend on overall likelihood and can be dominated by dense regions of the data, often favoring models that fit the bulk well but miss important features like tail behavior [55]. For example, take the case of the Mouse v117, where we fitted the model using *all data* rather than focusing on the tails (as in the other datasets; see Table 2). This introduces more influence from the bulk, especially at small values, which can affect the AIC/BIC values and explains why it differs from the other cases where tail-focused fitting reduces this effect. For all datasets, the selected models consistently show lower KS distances and better visual agreement with the data.

**Table 2:**
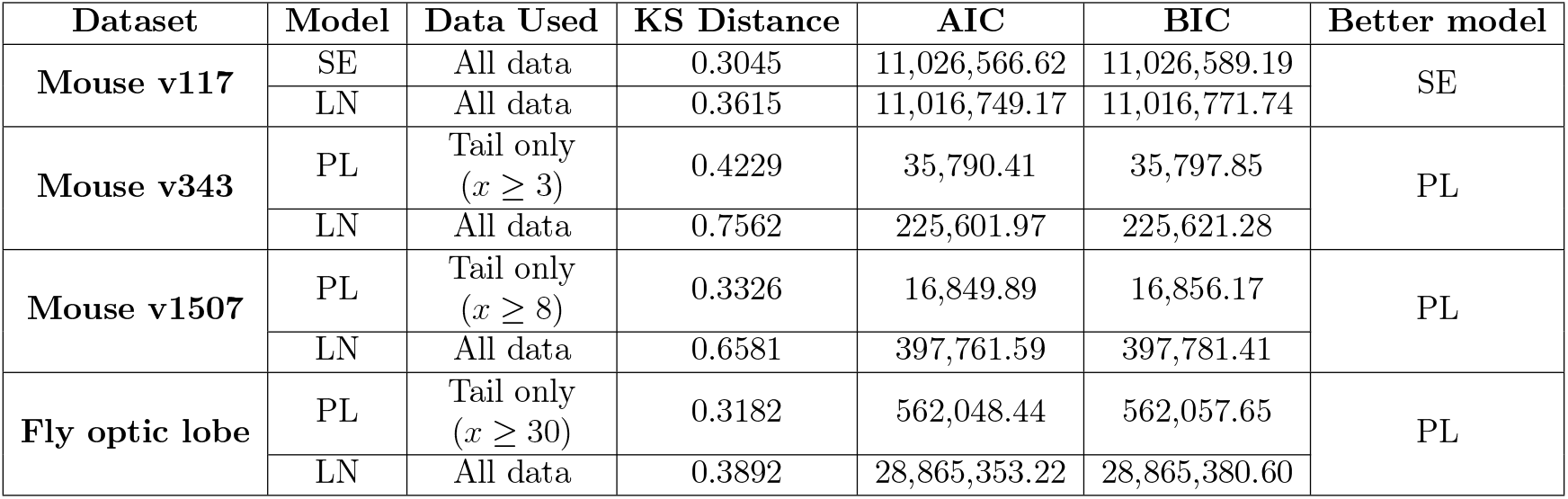
Comparison of statistical fits for synaptic edge weight distributions across datasets using KS distance, AIC and BIC methods. Lower values correspond to better fits and the better model is indicated in the last column.

| Dataset | Model | Data Used | KS Distance | AIC | BIC | Better model |
| --- | --- | --- | --- | --- | --- | --- |
| Mouse v117 | SE | All data | 0.3045 | 11,026,566.62 | 11,026,589.19 | SE |
|  | LN | All data | 0.3615 | 11,016,749.17 | 11,016,771.74 |  |
| Mouse v343 | PL | Tail only<br>($x \geq 3$) | 0.4229 | 35,790.41 | 35,797.85 | PL |
|  | LN | All data | 0.7562 | 225,601.97 | 225,621.28 |  |
| Mouse v1507 | PL | Tail only<br>($x \geq 8$) | 0.3326 | 16,849.89 | 16,856.17 | PL |
|  | LN | All data | 0.6581 | 397,761.59 | 397,781.41 |  |
| Fly optic lobe | PL | Tail only<br>($x \geq 30$) | 0.3182 | 562,048.44 | 562,057.65 | PL |
|  | LN | All data | 0.3892 | 28,865,353.22 | 28,865,380.60 |  |

**Table 3:**
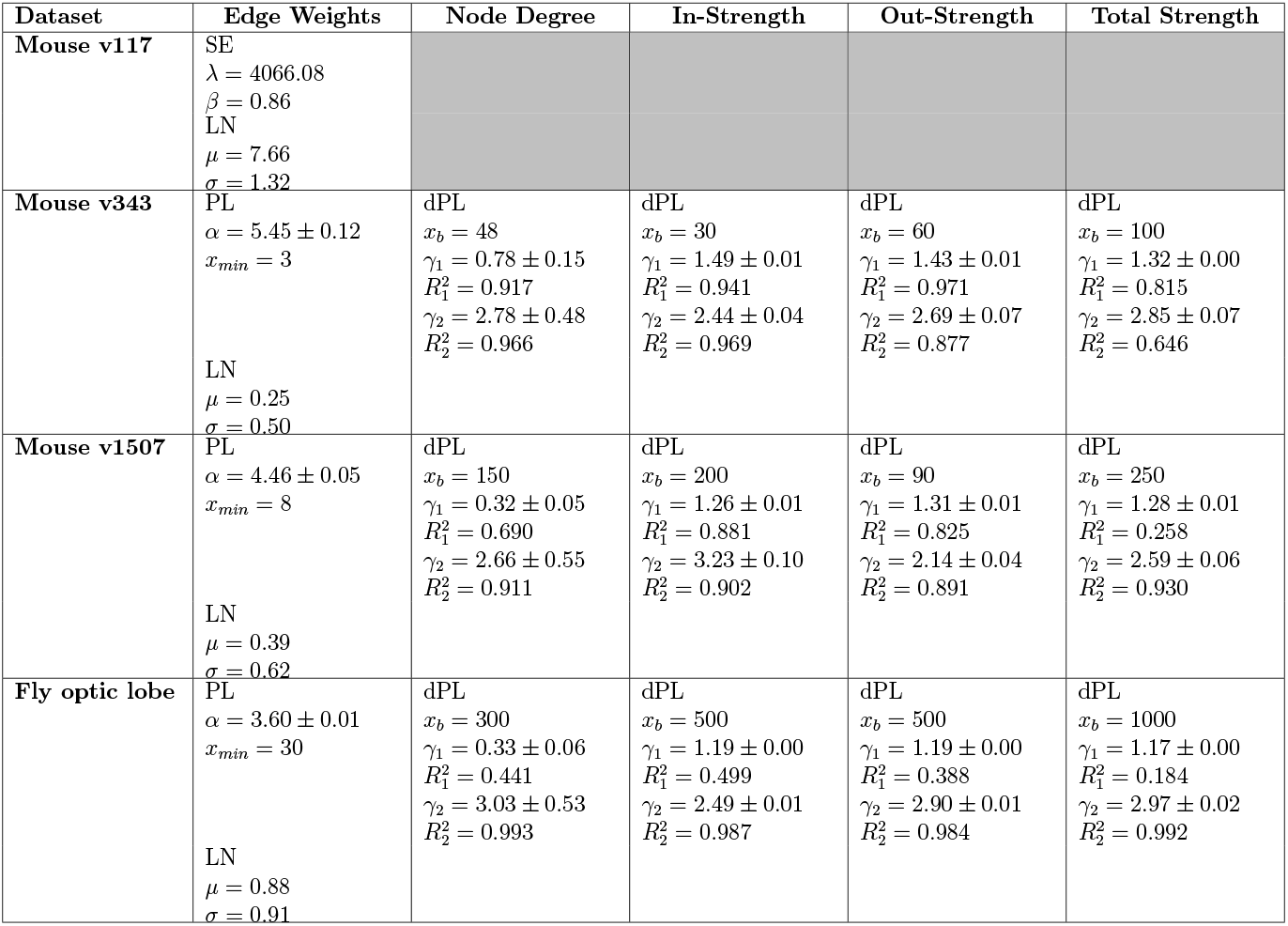
Distribution fits for edge weights, node degree, and node strengths (*s*^*in*^, *s*^*out*^, *s*^*tot*^) in the analyzed datasets. Reported parameters include breakpoints *x*_*b*_, scaling exponents *γ*_1_, *γ*_2_, and *R*^2^. Gray cells denote unavailable values.

In literature, a degree distribution that follows a PL means that the network is scale-free with few nodes that have very high number of connections, while most nodes have very few connections. Figure 2 shows that the degree distributions *P* (*k*) for all datasets in consideration follow a piecewise dual powerlaw (dPL) which may suggest two distinct organizational principles are at play. For other spatial networks such as airports, the dPL behavior observed in the degree distribution *P* (*k*) is attributed to a cost constraint in creating long-ranged links [56]. This constraint forces a node to abruptly reduce the length of new links once it reaches a critical degree *k*_*c*_. This sudden shift in growth strategy causes a change in the attachment probability, resulting in two different scaling behaviors: one for smaller nodes and one for major hubs above *k*_*c*_. This can also be true for brain networks as it is well-known that the brain also experiences such constraint in creating long links such that the distance distributions follow the exponential decay rule [57, 58, 59, 29].

**Figure 2:**
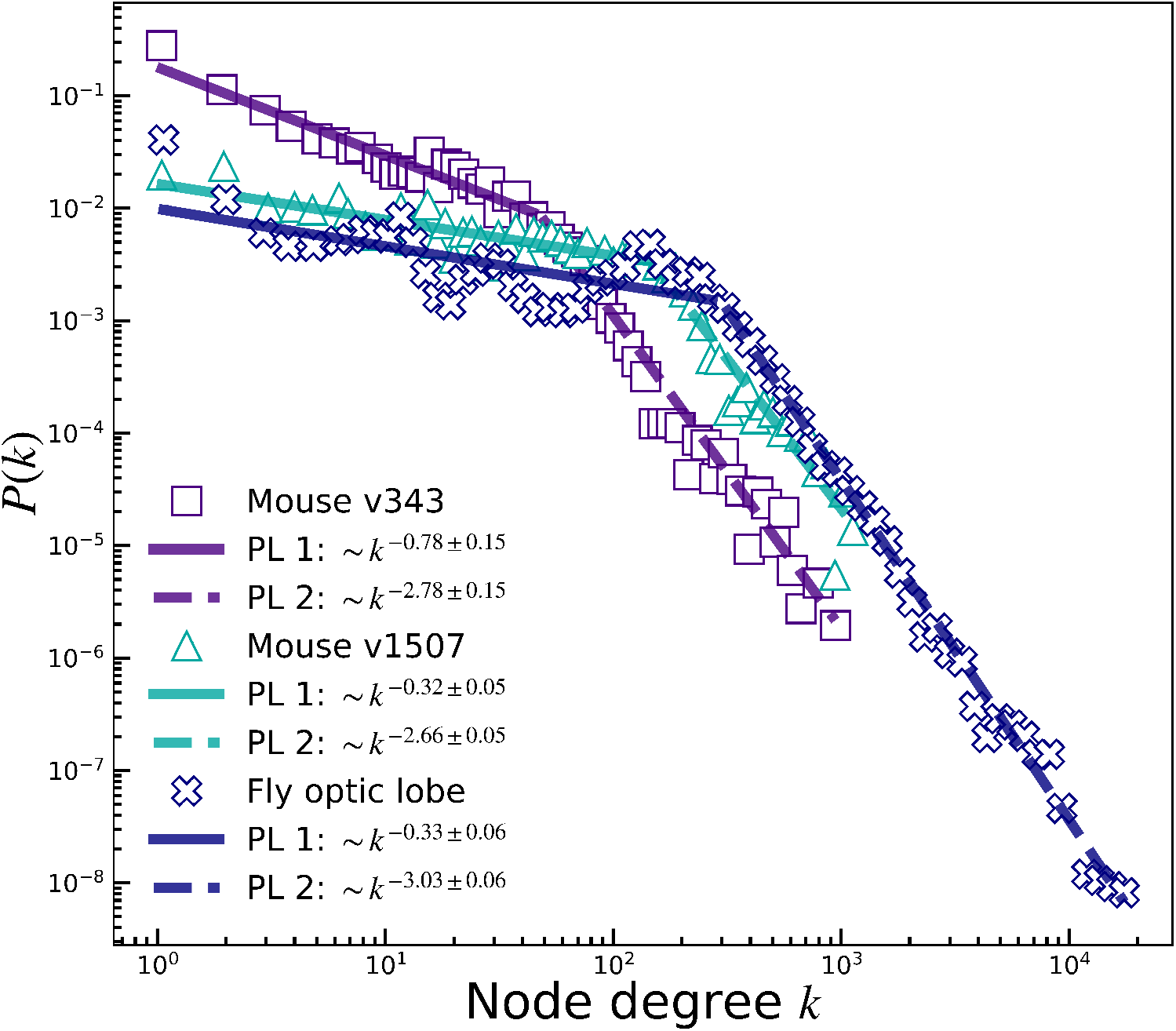
Node degree distributions. Node degrees of the network follow a dual power-law (dPL) distribution, highlighting heterogeneous connectivity. The distinct regimes may have also resulted from the spatial constraint of creating long-ranged links.

Figure 3 shows that the node strength distributions (*P* (*s*^*in*^),*P* (*s*^*out*^),*P* (*s*^*tot*^)) also behave according to a dPL behavior. We propose that the presence of the two PLs is indicative of different local topological contactome behavior, reflected by the dPL degree distributions. Furthermore, the crossover point shift signals the effect of a Hebbian learning on these structures. In a previous work [29], the local node strengths of various connectomes were found to follow a LN behavior. From a modelling standpoint, dual-scaling behavior has been a prominent feature in the edge weight distributions produced by self-organizing criticality (SOC) models [60, 61]. In Ref. [60], the initial, shallower power law (PL) tail was linked to short-range correlations within the base lattice, which diminish over time, while a steeper second PL tail emerges and strengthens as the learning process unfolds. Notably, in both our current study and prior research [29] which deal with empirical connectome data, we did not observe dual-power laws (dPL) in the global edge weight distributions. Instead, we found that node strengths (*s*^*in*^, *s*^*out*^, *s*^*tot*^) follow a LN distribution, and for the visual system connectomes, a dPL emerged.

**Figure 3:**
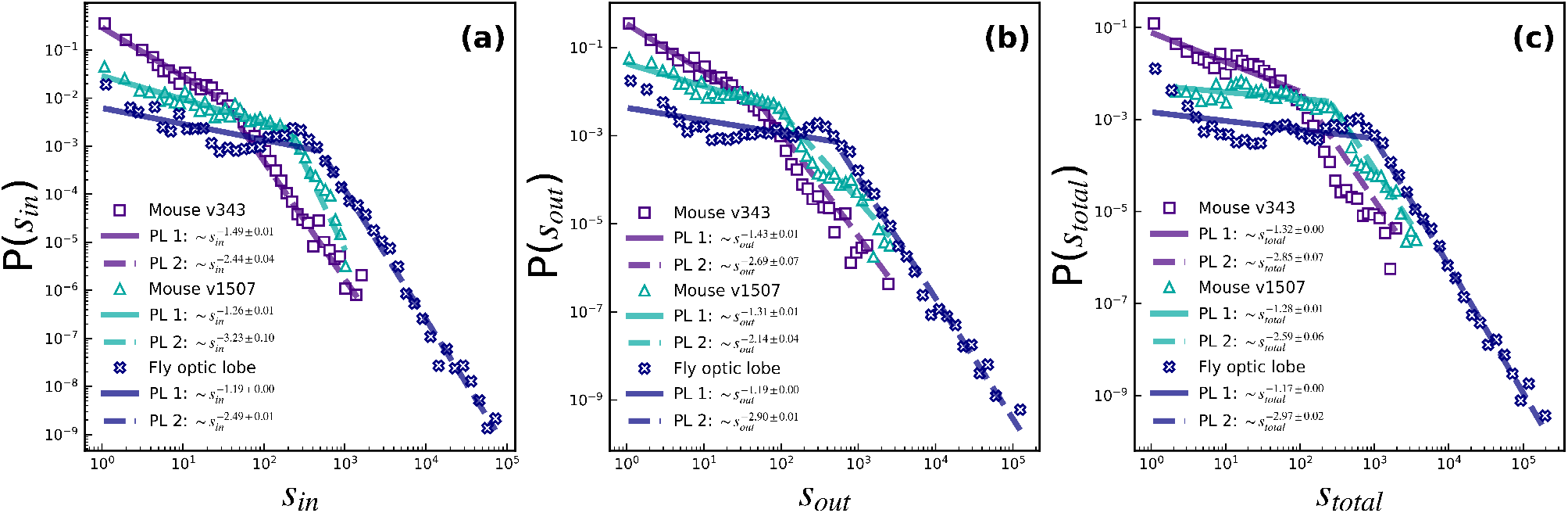
Local node strength distributions. Unlike previous findings on various connectomes, the visual connectomes in consideration are found to exhibit piecewise dPL node strengths, instead of LN.

So far, we have observed that the networks exhibit heterogeneous connectivity, with the node degrees following a dPL (Figure 2) and the node strengths, computed by summing the PL-distributed edge weights, also display dPL behavior (Figure 3). To investigate the possible origin of this observed dual scaling, we applied a null model analyses described in Section 2.

Figures 4(a)-(d) show the results for the random topology null models. Here, the *P* (*k*) and *P* (*s*) do not show dPL behavior. Since the nodes have characteristic number of connections, summing either uniform random or PL weights gives strengths that are mostly moderate. This tells us that random wiring, regardless of the added weights, cannot produce the dual scaling we observed in the data. In contrast, Figures 4(e)-(h) show the results for the degree-preserving null models, which keep the exact number of incoming and outgoing connections for each node. By doing so, these models preserve the heavy-tailed structure of the degrees (as shown in Figure 4(e)). Consequently, the computed node strengths (*s* : {*s*^*in*^, *s*^*out*^, *s*^*tot*^}) also exhibit dPL behavior for both weight-assignment schemes. However, with the uniform random weights, high-degree nodes add many moderate edge weights, producing very large *s* values. With PL weights, there is preponderance of low-edge weight values resulting in many nodes having lower strengths, shifting *s* to low-strength regime. These results show that the degree structure can be the key driver of the observed dual scaling in the node strengths distributions (*P* (*s*) : {*P* (*s*^*in*^), *P* (*s*^*out*^), *P* (*s*^*tot*^)}), and additional weakening or reinforcing weight heterogeneity, drawn from PL a distribution, modulates the strengths. In addition to this, results of another null model investigation is shown in Appendix A, where we start with an unweighted scale-free Barabasi-Albert (BA) network wiring and assign PL weights with different exponents. As one can observe, for *α* ≤ 1.5, corresponding to fatter tails than that of the degrees of the pure BA graph (*γ* = 3) the *P* (*s*) distributions develop a second regime. This simple illustration can be likened to a system that has an initial synthetic and reproducible heterogeneous structure as a substrate, and as the edge weights are reinforced, the distribution *P* (*s*) of the node strengths also evolves.

**Figure 4:**
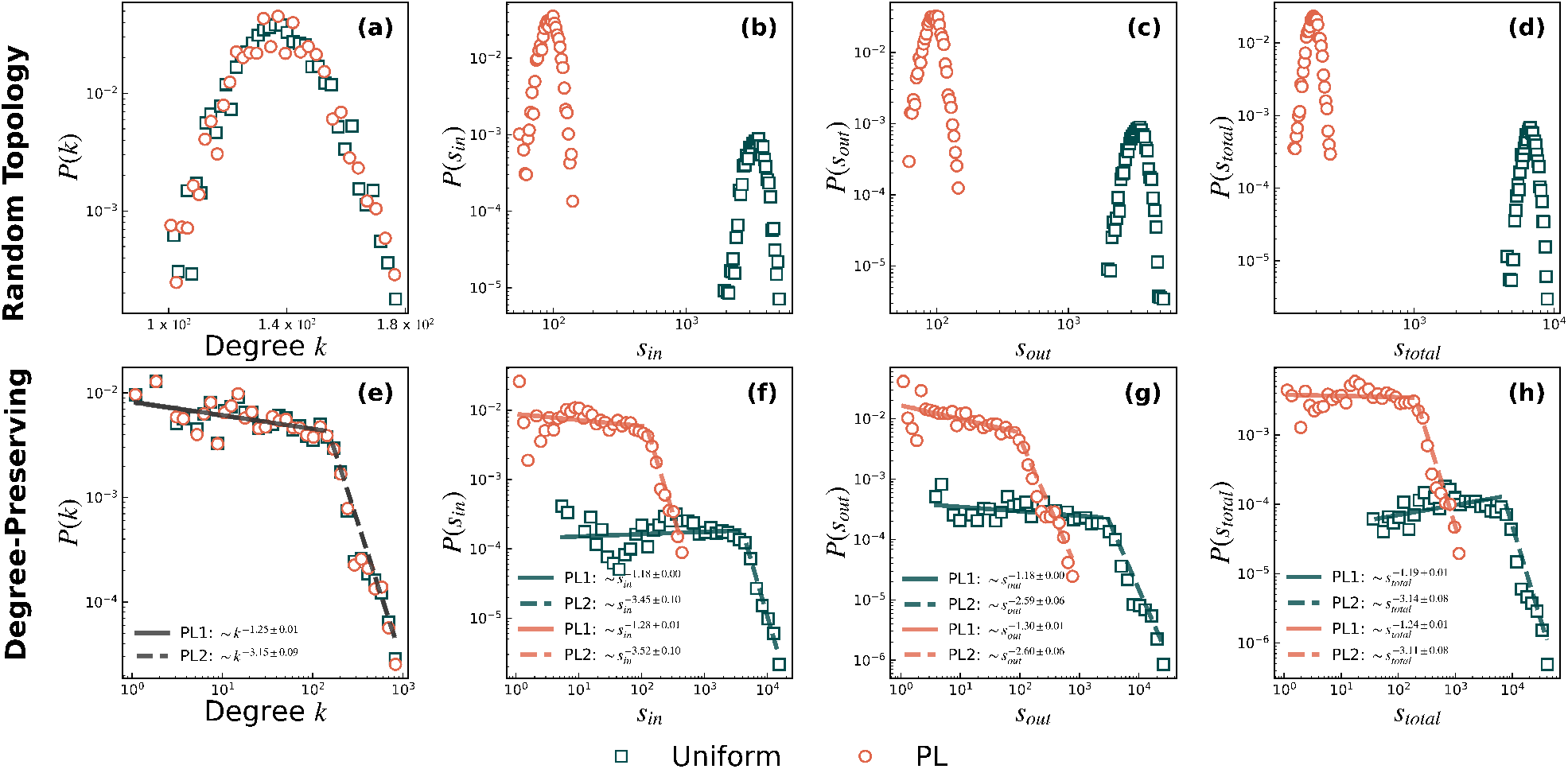
Degree and strength distributions for networks with random topology (a–d) and degree-preserving topology (e–h) (in the null-Models.) Columns show the distributions of degree (k), in-strength, out-strength, and total strength, respectively. For each case, results are shown for two edge-weight distributions: uniformly distributed weights (Random–uniform) (orange circles) and power-law (green squares) distributed weights. The degree-preserving null reproduces dPL behavior in both degrees and node strengths, whereas the random topology null fails to do so.

Having established that the node degree heterogeneity can provide dPL *P* (*s*) in the null models, we examine the contributions of the low-(80^*th*^ percentile) and high-(20^*th*^ percentile) degree *k* nodes to the node strengths of the real data, as shown in Figure 5. It turns out that low-*k* nodes mainly make up the first, lower-*s* regime, while high-*k* nodes dominate the heavy tail. However, when considering in-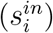 and out-strengths 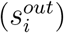 separately, the distinction between the contributions of the low- and high-*k* nodes becomes unclear. For example, some high-*k* nodes can appear in the low-*s* regime (as seen in Figs. 5(a),(b),(d),(e)) because not all their incoming and outgoing edges are strong. Similarly, a low-*k* node with a few particularly strong edges can sometimes have high 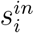 or 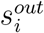.

**Figure 5:**
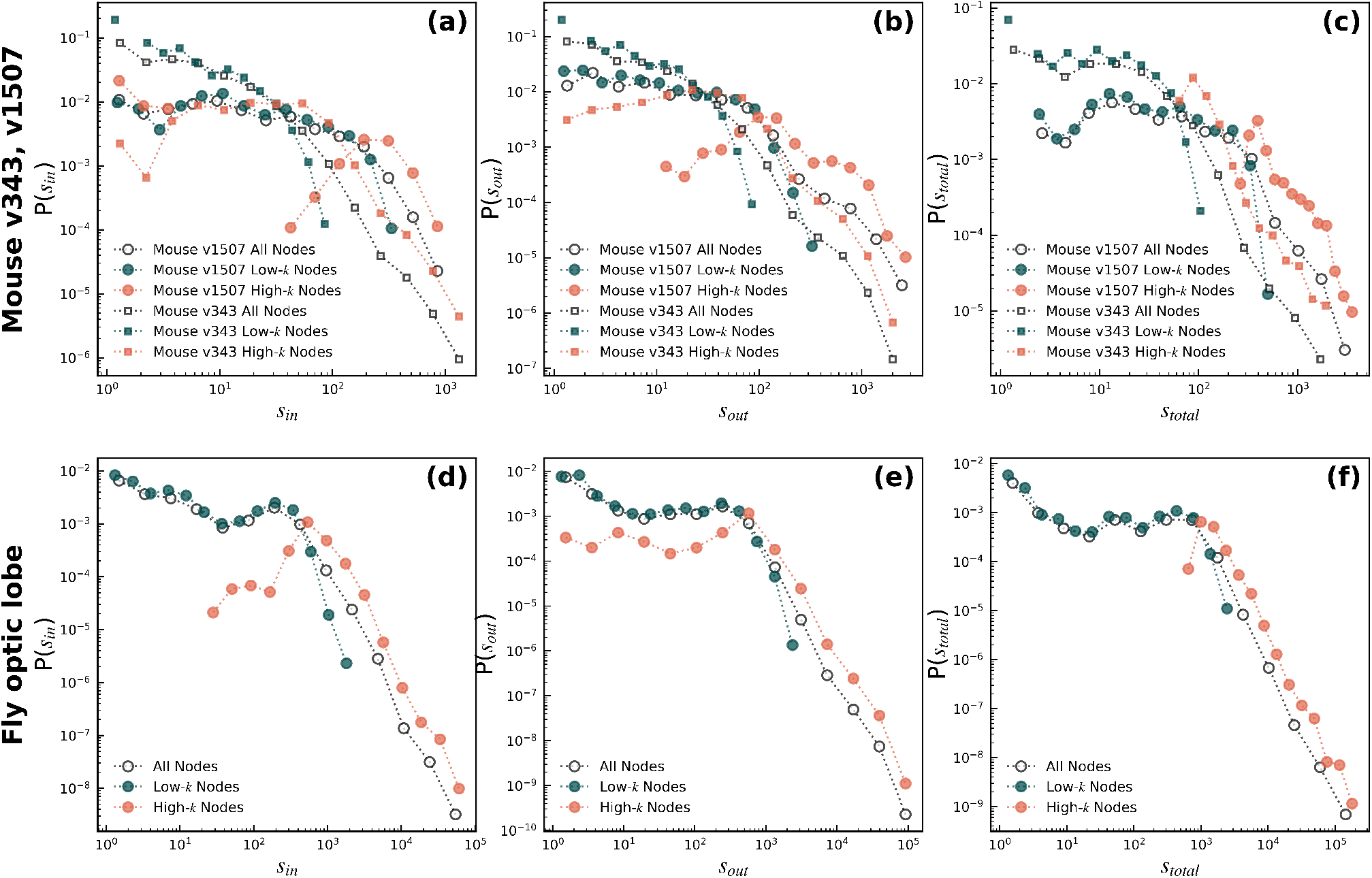
Contributions of low- and high-*k* nodes to node strengths *s*: low-*k* nodes form the first PL regime, while high-*k* nodes dominate the second (tail) regime, explaining the dual power-law structure.

By summing the 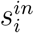 and 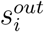 to compute for the total node strength *s*^*toti*^, these variations are averaged out, and the dPL becomes much more apparent (see Figs. 5(c) and (f)). In this view, it becomes clearer that the low-*k* nodes contribute the first PL regime, while high-*k* nodes dominate the second, heavy-tail regime. This demonstrates that the dPL behavior of the node strengths distributions *P* (*s*) is an emergent property from the combination of a heterogeneous connectivity and the distribution of edge weights across the network. In particular, the hubs (high-*k* nodes) drive the extreme tail matching we observed in the degree-preserving null model.

Having established that low- and high-*k* nodes contribute differently to the dPL behavior of the node strengths, we next examined how individual edges are organized with respect to node connectivity. The scale-free behavior of the degree distribution *P* (*k*) and the presence of PL behavior at the tails of the node strength distributions *P* (*s*) hint at the presence of hubs in the system characterized by high connectivity density. To probe this further, we analyzed the edge weights according to their directional profile allowing us to identify a group of segregable nodes that we dubbed *sources* and *sinks*. In the context of this work, source nodes are nodes that have purely outward connections 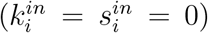, representing elements that strongly influence the network without being affected in return. Conversely, sink nodes have purely inward connections 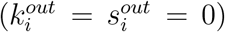, such that they integrate input from the network but exert little influence to others. These nodes represent neurons that can also be considered as *broadcasters* and *integrators*, defined and investigated in [62].

To gauge the influence of these source and sink nodes in the network, we quantify their relative abundance and the fraction of edges associated with them (Table 4). This provides a structural baseline for interpreting their role in the network.

**Table 4:** Percentage of source and sink nodes in the network and the share of incident edges.

| Dataset | % Nodes |  | % Incident Edges |  |  |
| --- | --- | --- | --- | --- | --- |
|  | Sources | Sinks | Sources | Sinks | Combined |
| Mouse v1507 | 1.38 | 0.78 | 0.91 | 0.11 | 1.01 |
| Fly optic lobe | 15.41 | 15.85 | 0.01 | 0.69 | 0.70 |

Although these nodes represent a limited portion of the network, their functional importance may not be reflected by their abundance alone, but rather how they are connected to others. It is therefore helpful to examine the distribution of edge weights *P* (*w*_*T*_) associated with source and sink nodes to understand how connectivity strength is organized around them (see Fig. 6).

**Figure 6:**
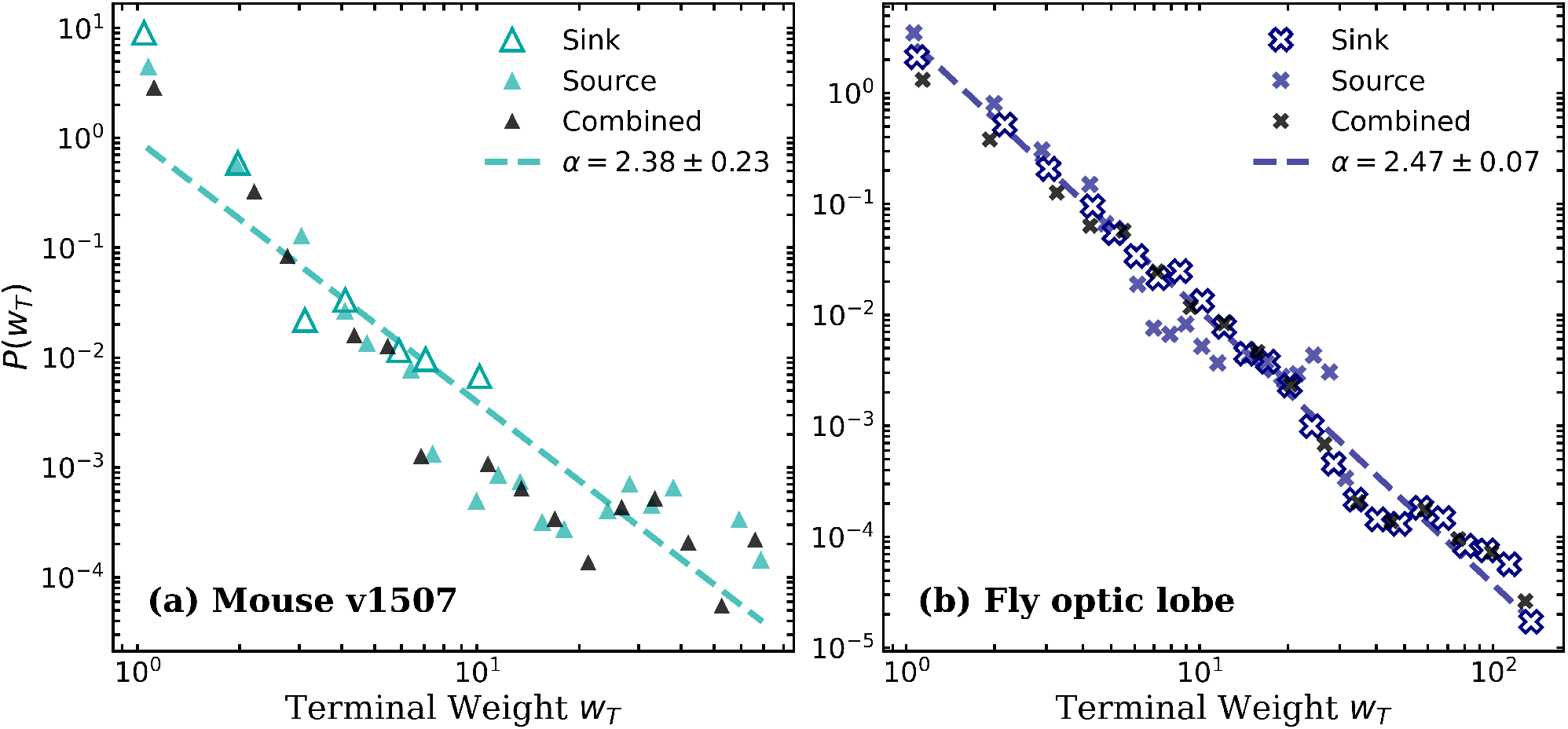
Edge weight distributions incident to the sources and sinks are scale-free, indicating that strong edges preferentially connect to highly connected nodes.

When we examine the weights of the edges incident to the source and sink nodes, we find that their behavior is best described by PL distributions: *P* (*w*_*T*_) ∼*w*^*−α*^ with exponents 2 *< α <* 3 in case of the most reliable connectomes (Mouse v1507 and the Fly optic lobe), spanning approximately two decades. This indicates clear scale-free behavior at the level of these directional subnetworks. Interestingly, prior work has shown that the global synaptic weight distributions for various connectomes follow PL tails with exponents *α* ≥ 3 [29]. In contrast, the smaller exponents observed here correspond to heavier tails, implying a higher probability of extremely strong connections around the source and sink nodes. Thus, while the connectomes exhibit relatively constrained global weight statistics, we observe strongly heterogeneous local structure around major information broadcasters and integrators. In particular, source and sink nodes support disproportionately strong connections, consistent with Hebbian mechanisms that amplify already influential nodes.

So far, these observations imply that strong connections are not random, but are instead linked to node connectivity (degree) and edge reinforcements that may occur depending on a node’s role in the network, as in the case of source and sink nodes. To make this relationship more explicit, we quantify the correlations between node degree, node strengths, and edge weights across the network. Specifically, we computed for the Pearson correlation coefficients among these quantities for various fractions of shuffled edge weights (Table 5; see Appendix B for details of the calculation procedure).

**Table 5:** Pearson correlation coefficients between node and edge properties as a function of the fraction of shuffled edges. Correlations are shown for *k*_*i*_ vs. *s*_*i*_ (*r*_*ks*_), *w*_*ij*_ vs. *K*_*ij*_ (*r*_*wk*_), and *w*_*ij*_ vs. *S*_*ij*_ (*r*_*ws*_). Unless otherwise noted, all correlations are statistically significant (*p* ≤ 0.05).

| Fraction shuffled<br>$f$ | Degree vs Strength ($r_{ks}$) | | Edge Weight vs Degree Sum ($r_{wk}$) | | Edge Weight vs Strength Sum ($r_{ws}$) | |
| --- | --- | --- | --- | --- | --- | --- |
|  | Mouse v1507 | Fly optic lobe | Mouse v1507 | Fly optic lobe | Mouse v1507 | Fly optic lobe |
| 0.00 | 0.973 | 0.874 | 0.249 | 0.065 | 0.276 | 0.108 |
| 0.05 | 0.975 | 0.881 | 0.238 | 0.061 | 0.262 | 0.102 |
| 0.10 | 0.976 | 0.891 | 0.226 | 0.058 | 0.249 | 0.095 |
| 0.15 | 0.978 | 0.898 | 0.209 | 0.054 | 0.232 | 0.089 |
| 0.20 | 0.980 | 0.906 | 0.202 | 0.052 | 0.222 | 0.084 |
| 0.25 | 0.982 | 0.914 | 0.187 | 0.048 | 0.206 | 0.077 |
| 0.30 | 0.982 | 0.923 | 0.177 | 0.044 | 0.196 | 0.071 |
| 0.35 | 0.985 | 0.931 | 0.159 | 0.042 | 0.176 | 0.066 |
| 0.40 | 0.986 | 0.939 | 0.150 | 0.037 | 0.165 | 0.058 |
| 0.45 | 0.988 | 0.948 | 0.135 | 0.035 | 0.148 | 0.053 |
| 0.50 | 0.989 | 0.953 | 0.123 | 0.031 | 0.136 | 0.048 |
| 0.55 | 0.991 | 0.961 | 0.115 | 0.029 | 0.126 | 0.042 |
| 0.60 | 0.992 | 0.968 | 0.097 | 0.025 | 0.108 | 0.036 |
| 0.65 | 0.992 | 0.974 | 0.087 | 0.023 | 0.097 | 0.032 |
| 0.70 | 0.994 | 0.980 | 0.071 | 0.019 | 0.080 | 0.026 |
| 0.75 | 0.994 | 0.983 | 0.062 | 0.017 | 0.070 | 0.023 |
| 0.80 | 0.995 | 0.989 | 0.049 | 0.012 | 0.056 | 0.016 |
| 0.85 | 0.996 | 0.992 | 0.039 | 0.009 | 0.045 | 0.012 |
| 0.90 | 0.996 | 0.995 | 0.026 | 0.006 | 0.032 | 0.008 |
| 0.95 | 0.997 | 0.996 | 0.012 | 0.004 | 0.018 | 0.005 |
| 1.00 | 0.996 | 0.997 | 0.000<br>$p = 0.911$ | 0.000<br>$p = 0.758$ | 0.007 | 0.002 |

In Table 5 we observe that as the shuffle fraction *f* increases, two opposing trends occur: (i) *r*_*ks*_ →1 indicating that under randomization a node’s strength is largely determined by its number of connections. In contrast, (ii) *r*_*wk*_ and *r*_*ws*_ →0, showing that these relationships depend on the specific placement of weights.

While node-level correlations (*r*_*ks*_) are strong, the correlations between edge weights and node properties (degree/strength), i.e., *r*_*wk*_ and *r*_*ws*_, are moderate but statistically significant (*p*≤ 0.05) at low shuffle fractions. This is expected in heterogeneous networks where most edges are weak, with only a few strong connections concentrated around key nodes (as seen in the PL-distributed edge weights in Figs. 1 and 6), which tend to co-activate. Biologically, edge weights in real connectomes reflect functional importance [41], wiring constraints [63], and possibly Hebbian-like reinforcement [41], as evidenced by the non-zero correlations (*r*_*wk*_, *r*_*ws*_) in the Mouse v1507 and Fly optic lobe at low *f*. Shuffling progressively disrupts this specificity, resulting in topology-driven networks with increasingly random weight placement. At full randomization (*f* = 1.0), correlations collapse to *r*_*wk*_ ≈ 0 and become statistically insignificant (*p >* 0.05), indicating that the association between weights *w*_*ij*_ and network structure *K*_*ij*_ has been effectively destroyed and is not expected under random assignment. Table 5 confirms these trends across both datasets, demonstrating that they are general rather than species-specific. Although the correlations are slightly stronger in the Mouse v1507 than in the Fly optic lobe network, the overall pattern remains consistent, emphasizing that network organization depends not only on degree and strength but also on the structured placement of edge weights.

We also analyzed the effect of shuffling on the *P* (*s*). To evaluate the influence of the structured organization of edge weights to the node strength distributions, we progressively shuffled increasing fractions (*f* ∈ [0, 1]) of edge weights while keeping the network topology fixed (Figure 7). This approach isolates the effect of weight placement from the degree structure and makes our results nontrivial. We found that small *f* causes the strongest deviations: the tail decays more rapidly, shifting away from its heavy-tailed behavior. As *f* increases, this effect weakens, and the tail becomes comparable to the original network.

**Figure 7:**
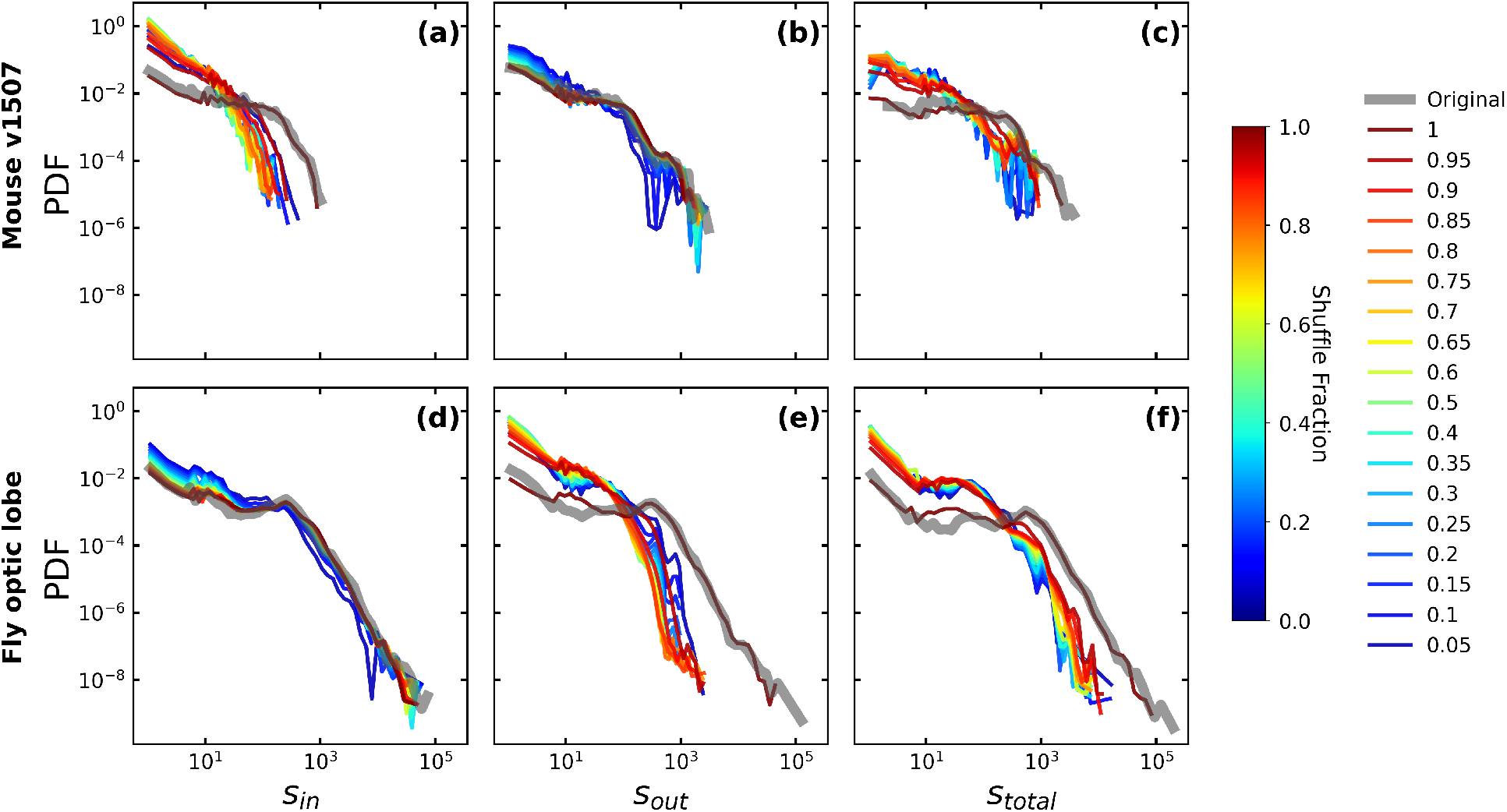
Effect of shuffling on the node strength distributions. in increasing fractions (*f* ∈ [0, 1]) shows that even small perturbations accelerate the decay of the heavy tail in node strengths, confirming the role of structured weight placement in amplifying the extreme-strength tail.

This pattern aligns with Table 5, where the correlation between edge weights and node properties (*r*_*wk*_, *r*_*ws*_) decreases as the shuffle fraction increases, while the correlation between node degree and total strength (*r*_*ks*_) remains high. These results suggest that the heavy tails in node strengths *P* (*s*) depend on the alignment of edge weights with node connectivity, and disrupting this alignment reduces the heaviness of tails of the node strength distribution *P* (*s*) without affecting the proportionality between *k* and *s*. Notably, shuffling has different effects on the mouse and fly data: it impacts in-strength (*s*^*in*^) more in the mouse, and out-strength (*s*^*out*^) more in the fly, reflecting structural differences between their connectomes.

To summarize, these results suggest that the dPL behavior in node strength distributions arises from two combined effects: the structure of the network (degree distribution *P* (*k*)) sets the baseline dual scaling. Second, the structured placement of strong edges, akin to Hebbian-like reinforcement, enhances the heavy tail. This layered organization demonstrates how topology and weight correlations interact to shape the observed network properties.

### Discussion on the difference between node strength and global synaptic weight distributions

As mentioned in Section 1, it is generally known that heavy-tailed statistical distributions are produced by many features of brain dynamics. At present, there is an ongoing debate on the contradicting observations of LN and PL behavior of the node strengths *P* (*s*) and edge weights distributions *P* (*w*), respectively, which may be related to critical behavior. In the following, we propose a simple explanation for the deviation of *P* (*s*) from the global PL behavior of *P* (*w*).

Node strengths can be regarded as generalized node degrees, when we ignore the identities of the connected nodes. They are simply the sums of incoming and outgoing edge weights for each node. Let’s assume, based on our criticality + Hebbian learning working hypothesis [29], that the edge/synaptic weights follow a global PL distribution (or at least have a PL tail).

For PL distributions with *α* > 3, the sum of independent and identically distributed random variables tends to a normal (Gaussian) distribution, just like in standard, non-heavy-tailed cases. This happens because when the variance of the variables is finite, the Central Limit Theorem (CLT) applies. This property may help explain why global and local weight distributions often differ. However, the assumption of independence of weight variables is only approximate. We performed a correlation analysis, by calculating ⟨|*w*_*i,j*_ −*w*_*ik*_|⟩ _*i*_*/* ⟨*s*_*i*_⟩ _*i*_, which resulted in less than 1% value, both for the Mouse (v1507) and the Fly connectomes (more details in the Appendix). This means that practically we can neglect the correlations and apply the CLT. In reality, due to this omissions the resulting *P* (*s*) distributions may not be perfectly Gaussian, but somewhat heavier-tailed, which can explain the observed lognormal distribution in many of the cases discussed in [29]. Here, we disregard dPL results, as they likely arise from specific structural features of the dPL contactome.

If *α* ≤3, the standard CLT does not apply. Instead, the sum of random variables in this type of distribution will follow a Lévy stable distribution, not to a Gaussian. The Generalized Central Limit Theorem (GCLT) explains this behavior, specifically for distributions with infinite variance. Lévy stable distributions have fat tails that drop off according to a PL, similar to the PL behavior seen in real-world networks. In Figure 6, the distributions of the weights of the edges incident to the source and sink nodes of our most accurate Mouse (v1507) and Fly optic lobe connectomes follow a scale-free pattern, which aligns with the GCLT prediction of PL tails for node strength distributions (*P* (*s*)), granted that there are enough sink/source nodes in the system (see Table 4).

Note, however that having not enough data can also introduce finite size cutoff in PL-s, thus many of former analysis claiming LN distributions should be considered with some criticism

## 4. Conclusion

In this study, we compare the topological organization of the male *Drosophila melanogaster* optic lobe and the mouse visual cortex. In summary, both datasets exhibit global synaptic weight distributions *P* (*w*) following PL behavior with the degree of connectome proofreading drastically influencing the apparent heaviness of distribution tails, with fully proofread datasets enabling more reliable identification of both weak and strong nodes. Additionally, our analyses showed that both datasets exhibit dPL distribution of local node strengths *P* (*s*), with low-*k* nodes contributing to the first scaling regime and high-*k* nodes forming the heavy-tailed second regime. Degree-preserving null models reproduced this pattern, whereas Random topology nulls do not, indicating that the observed dual-scaling is primarily determined by the underlying heterogeneous degree distribution of the network contactome structure, which is further modulated by the weights. Additionally, the examination of edge weights around source and sink nodes showed that strong connections are concentrated around high-*k* and/or key nodes that serve as broadcasters and integrators, following scale-free distributions 2 *< α* ≤ 3 suggesting local reinforcement mechanisms strengthened edges associated with key nodes. On the other hand, the correlation analyses (Table 5) and edge weight-shuffling (Figure 7) confirm that the heavy-tails of the node strength distributions *P* (*s*) depend on the specific alignment of edge weights and node connectivity (degree) such that disrupting this alignment by shuffling reduces the heaviness of tail behavior while leaving the degree-strength proportionality largely intact. Apart from these, the differences in directional profiles between the species i.e. mouse in-strength *P* (*s*^*in*^) and the fly’s out-strength *P* (*s*^*out*^) behavior under shuffling, highlight that the reinforcements of edges depend on the network’s wiring patterns. Moreover, to reconcile previously conflicting reports of PL and LN statistics in neural systems, we propose a mechanistic explanation in which distinct organizational scales give rise to different distributional regimes. Collectively, these findings indicate that reinforcement mechanisms shapes network structure and modulates weights across the system. Such organization, coupled with the sensitivity of the network to even partial randomization, is consistent with features of near-critical brain networks maintaining balance between global integration and local dynamics for optimal information processing.

## CRediT authorship contribution statement

**M.T.C**. did the data querying and fitting with empirical data and tested the PL fits; **I.P**. did the data querying and also some statistical tests; **G. Ó**. conceptualized the work and did the theoretical analysis; All authors analyzed the results, wrote the draft, and reviewed the manuscript.

## Competing interests

The authors declare no known competing financial or personal interests.

## Data Availability

Data is available upon request from G. Ódor via email at.

## Research Funding

This research was conducted with support from the Hungarian National Research, Development and Innovation Office NKFIH(K146736).

## Appendix A. Sparse BA with the addition of varying *α* weight exponents

**Figure A8:**
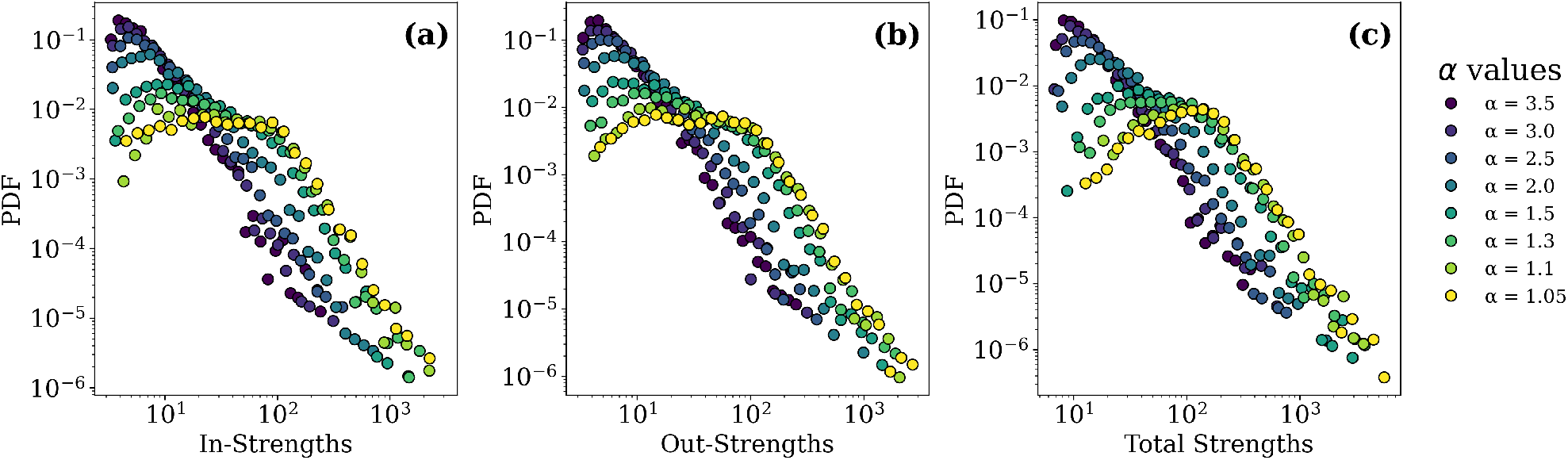
A sparse BA network. (*N* = 2178, *m* = 3, *E* = 13, 050) structure, with additional weights drawn from PL distribution of heavier tails (low exponent values: i.e. *α* ≤ 1.5) can create a graph with dPL behavior in the in-/outstrength distributions.

## Appendix B. Node and Edge-level Pearson Correlations

**Figure B9:**
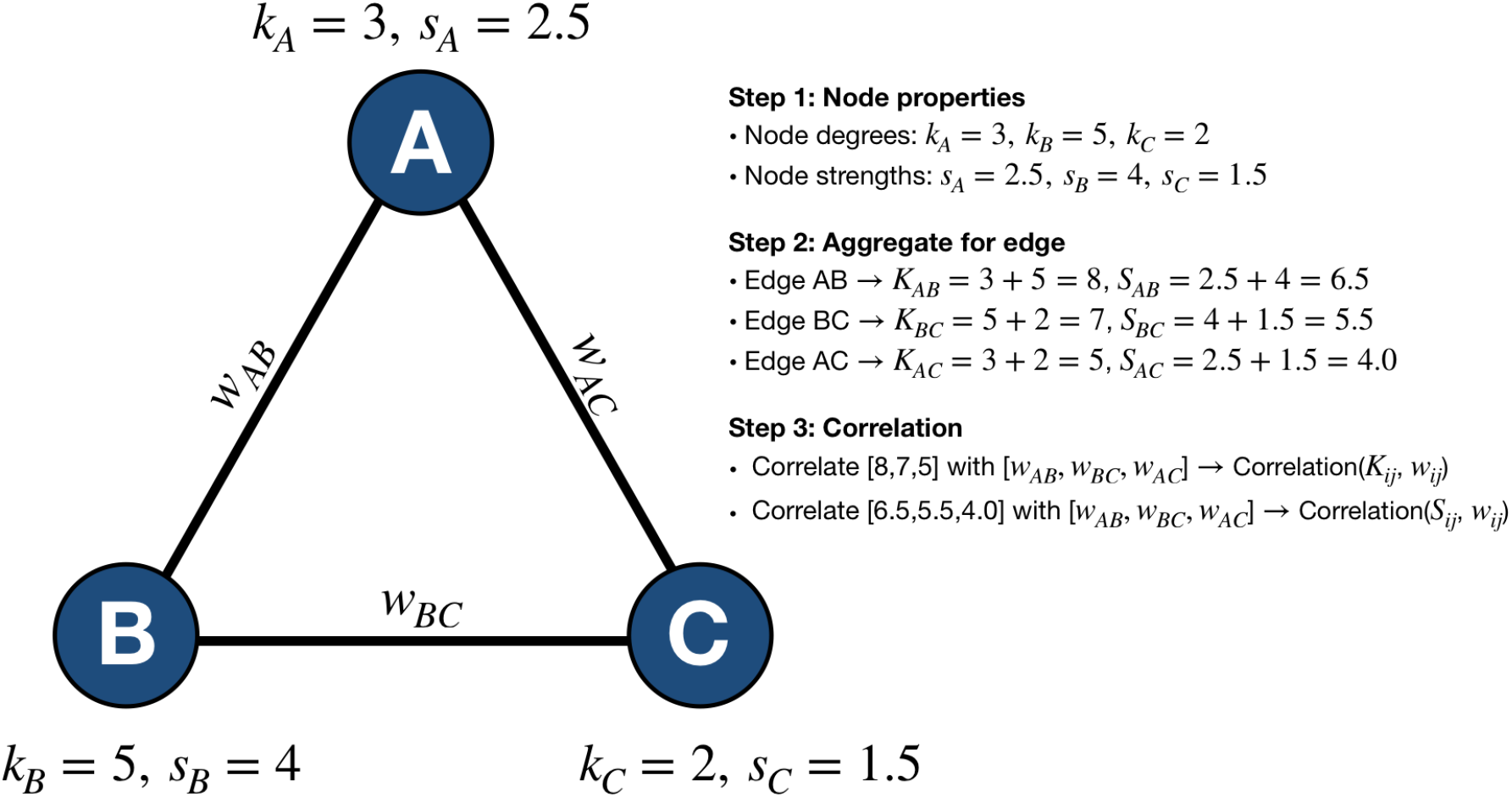
Schematic illustration of node and edge-level correlations in a weighted, undirected network. Each node *i* has an associated degree (*k*_*i*_) and strength (*s*_*i*_), the sum of edge weights). For each edge (*i, j*) with weight *w*_*ij*_, the sums of its endpoint degrees (*K*_*ij*_ = *k*_*i*_ + *k*_*j*_) and strengths (*S*_*ij*_ = *s*_*i*_ + *s*_*j*_) are computed. These aggregated quantities are then correlated with the edge weights to quantify how node-level properties relate to edge weights.

To evaluate the extent of the specific assignment of edge weights contributes to the observed network properties, specifically the node strength distributions, we implemented a weight-shuffling null model on the empirical networks. Starting from the original directed weighted graph, we systematically randomized edge weights while preserving the underlying topology.

Specifically, for a given fraction (*f* ∈ [0, 1]), a subset of edges corresponding to *f* of the total edge set was selected uniformly at random, and their weights were permuted among themselves without replacement. This procedure maintains the global weight distribution and network connectivity but progressively disrupts the specific assignment of weights to edges.

For each level of randomization, node strengths *s*_*i*_ were recomputed, while node degrees *k*_*i*_ remained unchanged. We then quantified three Pearson correlation relationships (with confidence levels characterized by *p*-values):

i. *r*_*ks*_, for *k*_*i*_ vs *s*_*i*_,
ii. *r*_*wk*_, for *w*_*ij*_ vs *K*_*ij*_,
iii. *r*_*ws*_, for *w*_*ij*_ vs *S*_*ij*_ .

For cases (ii) and (iii), the node properties, *k* and *s* were represented by the sum of the source and target node values, yielding a single value (*K*_*ij*_, *S*_*ij*_) that captures the joint contribution of both nodes to each edge. These metrics capture the coupling between network topology and weight organization at both node and edge levels. The analysis was repeated across a range of shuffle fractions *f* ∈ [0, 1] with increments of: Δ*f* = 0.05, with independent random seeds for each realization. All computations were implemented in Python using the igraph library, and parallelized across shuffle levels to improve computational efficiency.

## Appendix C. Correlation among incoming weights of nodes

For each node *i*, we quantify how heterogeneous its incoming connection weights are by employing the following steps. First, we extract the incoming weights {*w*_*ji*_ |*w*_*ji*_ *>* 0} and compute for all pairwise absolute differences |*w*_*ji*_ −*w*_*li*_| then we take the mean of these differences for every node.

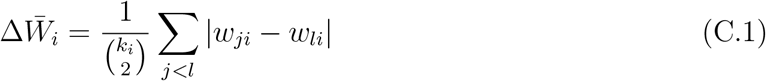

where *k*_*i*_ is the number of incoming edges to node *i*, and *w*_*ji*_, *w*_*li*_ are incoming weights.

To assess whether 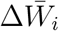 is structurally driven, we examined the relationships between heterogeneity 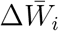 and the number of incoming connections *k*_*i*_ and the total incoming weight *s*_*i*_.

**Table C6:** Summary of average within-node edge weight heterogeneity (*< H*_*i*_ *>*), average number of incoming connections 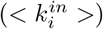, and total incoming weight 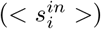 for nodes across different datasets.

| Datasets | $\langle \Delta \bar{W}_i \rangle$ | $\langle k_i^{in} \rangle$ | $\langle s_i^{in} \rangle$ |
| --- | --- | --- | --- |
| Mouse v1507 | 1.270 | 70.045 | 134.703 |
| Fly optic lobe | 4.907 | 118.950 | 487.340 |

## Supplementary Material

### Appendix C.1. Statistical distributions

In this work, we employed logarithmic binning for both datasets and fitted them with the following distributions. Below are their mathematical descriptions and parameters.

### 1. Exponential Distribution

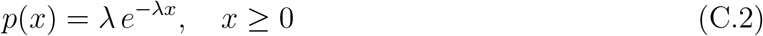

where *λ >* 0 is the rate parameter

### 2. Stretched Exponential Distribution

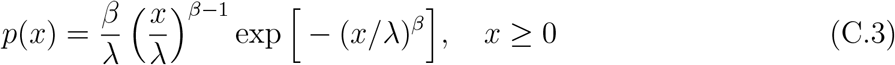

where *λ >* 0 is the scale parameter, *β >* 0 is the shape parameter

### 3. Lognormal Distribution

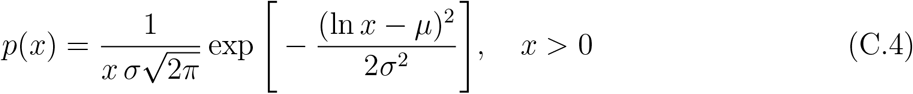

where *µ* and *σ >* 0 are the mean and standard deviation of ln *X*

### 4. Power Law

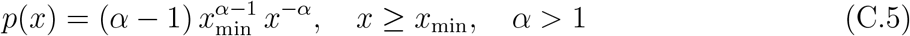

*α* is the scaling exponent, *x*_*m*_*in* is the lower cutoff.

### 5. Piecewise Dual Power Law

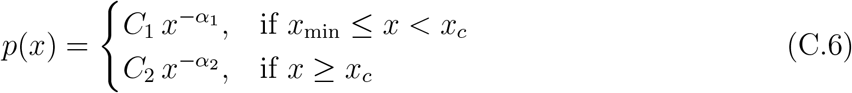

where the normalization constants *C*_1_ and *C*_2_ are defined as follows,

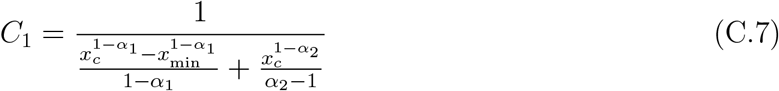

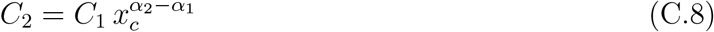

where *α*_1_, *α*_2_ are the two exponents, *x*_*c*_ is the crossover point, *x*_*m*_*in* is the lower cutoff. This ensures continuity at *x* = *x*_*c*_ and that the total probability integrates to 1.

### Appendix C.2. Powerlaw exponent estimation

To characterize the heavy-tailed nature of the weight distributions, we fitted a power-law (PL) model to the tail of the data using a standard maximum likelihood estimator (MLE). For continuous data with support *x* ≥ *x*_min_, the PL probability density function is

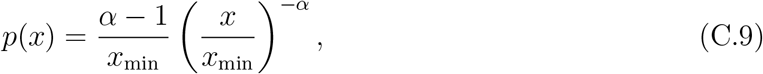

where *α >* 1 is the scaling exponent and *x*_min_ is the lower bound of the scaling region.

Following the approach of Clauset et al. (2009) [**?**], we estimated *α* from the subset of data satisfying *x* ≥ *x*_min_ by maximizing the log-likelihood function

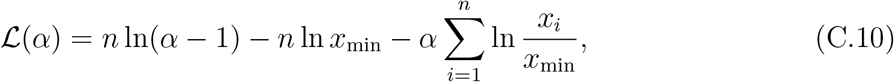

where *n* is the number of data points in the tail.

The MLE solution for the scaling exponent is

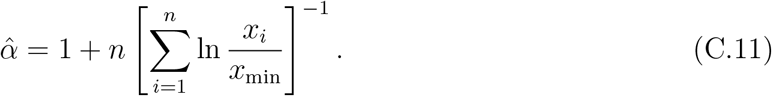

This estimator is unbiased in the large-sample limit and is commonly used for fitting PL tails in empirical distributions.

To quantify the uncertainty of the estimate, we used the standard error of the MLE, derived from the Fisher information of the Pareto distribution:

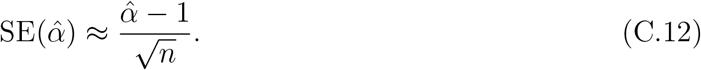

### Appendix C.3. Determining the Better Model for Edge Weight Distributions

When comparing statistical fits for edge weight distributions, three main metrics are used: Kolmogorov-Smirnov (KS) distance [64], Akaike Information Criterion (AIC) [65, 66], and Bayesian Information Criterion (BIC) [67]. The *better model* column in the table reflects the fit that is most consistent with the data according to these metrics.

The **Kolmogorov-Smirnov (KS) distance** measures the maximum difference between the empirical cumulative distribution function (CDF) of the data and the CDF of the fitted model. A smaller KS value indicates that the model closely follows the observed distribution. Its continuous form is given by

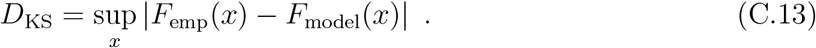

On the other hand, for discrete data (with sample of size *n*) takes the form,

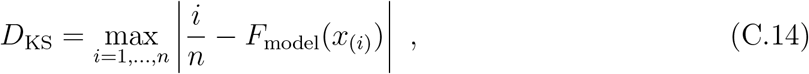

where *F*_*emp*_(*x*) is the empirical CDF of the data, *F*_*model*_(*x*) is the model CDF, *x*_*i*_ are the datapoints, and *n* is the sample size.

The **Akaike Information Criterion (AIC)** estimates the relative quality of a statistical model by balancing goodness-of-fit and model complexity:

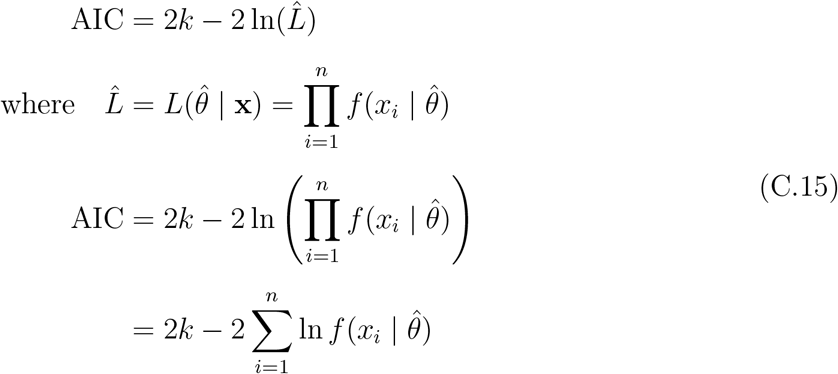

where *k* is the number of parameters, 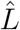 is the maximum likelihood of the model given the data, and *f* (*x*_*i*_ |*θ*) is the probability density (or mass) function of observables *x*_*i*_ given parameters *θ*. A lower AIC indicates a better balance of fit and simplicity. For example, for Mouse v117, stretched exponential’s (SE) AIC is slightly higher than LN, but visually and via KS, SE better captures the distribution, indicating that AIC alone is not always the deciding factor when distributions are heavy-tailed or data are sparse.

The **Bayesian Information Criterion (BIC)** is similar to AIC but penalizes model complexity more strongly:

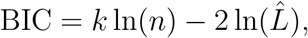

where *n* is the number of observations. Again, a lower BIC suggests a better model and is particularly useful when comparing models with different numbers of parameters.

## Notes

### Competing Interest Statement

The authors have declared no competing interest.

### Summary of Updates

This version of the manuscript is a major overhaul of the first one containing more analytical and numerical model calculations to support the structural connectome distributions. In particular it presents comparisons with null models and correlation analyses, which suggest that the underlying contactome structures exhibit double power-law degree distributions, modified by weights, in agreement with critical Hebbian learning mechanism. The present version is accepted in Chaos&Solitons&Fractals.

